# Widespread mermithid nematode parasitism of Cretaceous insects

**DOI:** 10.1101/2023.02.07.527443

**Authors:** Cihang Luo, George O. Poinar, Chunpeng Xu, De Zhuo, Edmund A. Jarzembowski, Bo Wang

## Abstract

Mermithid nematodes are obligate invertebrate parasites dating back to the Early Cretaceous. Their fossil record is sparse, especially before the Cenozoic, thus little is known about their early host associations. This study reports 16 new mermithids associated with their insect hosts from mid-Cretaceous Kachin amber, 12 of which include previously unknown hosts. These fossils indicate that mermithid parasitism of invertebrates was already widespread and played an important role in the mid-Cretaceous terrestrial ecosystem. Remarkably, three hosts (bristletails, barklice and perforissid planthoppers) were previously unknown to be parasitized by mermithids both past and present. Furthermore, our statistical analyses show that in contrast to their Cenozoic counterparts, Cretaceous nematodes including mermithids are more abundant in heterometabolous insect hosts. This result suggests that nematodes have not completely exploited the dominant Holometabola as their hosts until the Cenozoic. This study reveals what appears to be a vanished history of nematodes that parasitized Cretaceous insects.

## Introduction

Nematodes (roundworms), a group of non-segmented worm-like invertebrates, are one of the most abundant animals on earth in terms of individuals (*Lorenzen, 1994; Poinar, 2011*). They are distributed worldwide in almost all habitats and play key roles in ecosystems by linking soil food webs, influencing plant growth and facilitating nutrient cycling (*Yeates et al., 2009; van den Hoogen et al., 2019; Zhang et al., 2020*). The earliest known definite nematode fossils occur in the Lower Devonian Rhynie Chert inside the cortex cells of an early land plant and have been considered to be a plant parasite (*Poinar et al., 2008*), but unconfirmed nematode-like fossils and trace fossils can date back to the Precambrian (*Poinar, 1979*). Despite their abundance in many extant ecosystems, nematodes are exceedingly rare in the fossil record, since most of them are small with soft bodies and concealed habits (*Poinar, 2011*).

The Mermithidae represent a family of nematodes that are obligate invertebrate parasites which occur in insects, millipedes, crustaceans, spiders, molluscs and earthworms (*Nickle, 1972; Poinar, 1979*). They can affect the morphology, physiology, and even the behaviour of their hosts (*Petersen, 1985*). The life cycle of mermithids comprises five stages (*Poinar and Otieno, 1974; Poinar, 1983*, *2001b*). Eggs are deposited in the environment, and the developing embryo moults once and emerges from the egg as a second-stage juvenile. This juvenile is the infective stage that enters the hemocoel of a potential host. After a relatively rapid growth phase (third stage), the mermithid exits the host as a postparasitic juvenile (fourth stage). It then enters a quiescent phase and moults twice to become an adult (*Poinar, 2015*). Hosts usually die when the mermthids exit, which is why mermithids have been widely studied as possible biological control agents, especially against aquatic stages of medically important insects like mosquito larvae (*Petersen, 1985*). Although mermithids kill their hosts like parasitoids, they are commonly considered as parasites like other nematodes *(De Baets et al., 2021)*.

Paleontological records of parasitism provide critical clues about the origination and diversification of important nematode groups, and elucidate the synecology and coevolution of ancient parasites and their hosts. Due to their large size and invertebrate parasitic habits, mermithid nematodes are most likely of all nematodes to occur as recognizable fossils, especially in amber as they exit their invertebrate hosts (*Poinar, 2011*). Their fossil record dates back to the Early Cretaceous with *Cretacimermis libani* from a chironomid midge (Diptera: Chironomidae) in Lebanese amber (~135 Ma; million years ago) (*Poinar et al., 1994*). However, there is a dearth of examples of other Cretaceous mermithids prior to the present study, with only *Cretacimermis chironomae*, also from chironomid hosts, *Cretacimermis protus* from biting midges (Diptera: Ceratopogonidae) and *Cretacimermis aphidophilus* from an aphid (Hemiptera: Burmitaphididae) in mid-Cretaceous Kachin amber (*Poinar and Buckley, 2006; Poinar and Sarto i Monteys, 2008; Poinar, 2011*, *2017*).

Here, we report 16 additional mermithid nematodes associated with their insect hosts in mid-Cretaceous Kachin amber (approximately 99 million years old). These examples triple the diversity of Cretaceous mermithids (from 4 to 13 species) and reveal previously unknown host–parasite relationships. Our study also shows that mermithids were widely distributed in a number of diverse insect lineages by the mid-Cretaceous, thus providing novel insights into the early evolution of nematode parasitism of insects.

## Results

### Systematic palaeontology

Due to the inability to adequately detect all biological adult characters, it is impossible to place or refer fossil mermithid nematodes to any natural extant genus. This is why fossil collective genera have been erected under the same guidelines as recent collective genera for difficult nematodes. The importance of placing nematode species in collective genera is to underpin or establish the time, place and hosts of these parasitic lineages. For Cretaceous Mermithidae not assignable to any previously known genus or lacking biologically preferred diagnostic characters, the collective genus *Cretacimermis* was erected (*Poinar, 2001b*). Putative hosts were determined by noting nematodes emerging from their bodies or completely emerged nematodes adjacent to potential hosts, especially if there is physical evidence that a particular insect was parasitized.

**Family** Mermithidae Braun, 1883

**Collective genus** *Cretacimermis* Poinar, 2001

**Collective species** *Cretacimermis incredibilis* Luo & Poinar, sp. nov. (Figures 1A and 2A–D)

**Figure 1.**
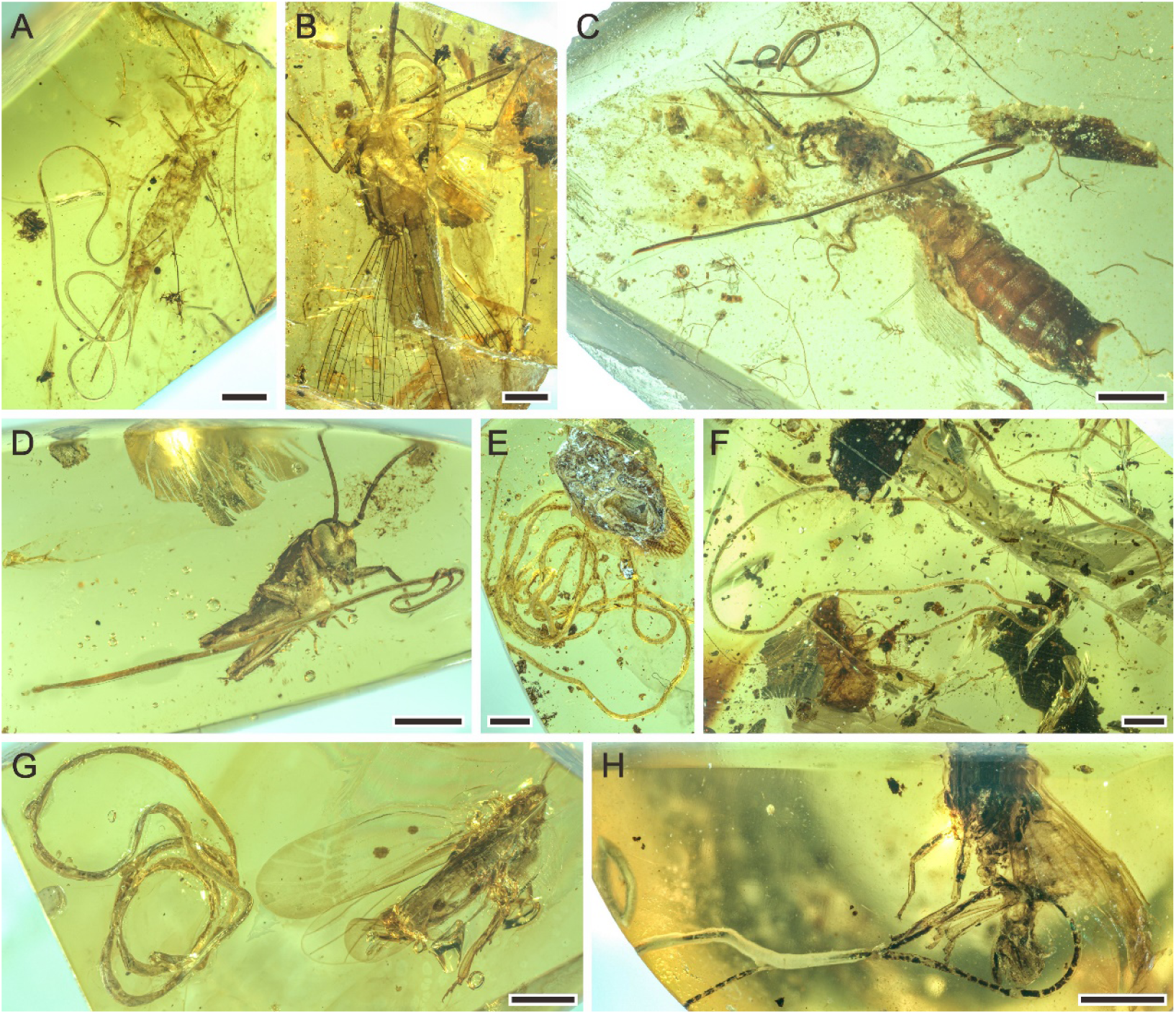
Mermithids and their insect hosts from mid-Cretaceous Kachin amber (~99 Ma; million years ago). Part I. (**A**) *Cretacimermis incredibilis* sp. nov. adjacent to its bristletail host. (**B**) *Cretacimermis calypta* sp. nov. adjacent to its damselfly host. (**C**) two separate specimens of *Cretacimermis adelphe* sp. nov. that have emerged from their earwig host. (**D**) *Cretacimermis directa* sp. nov. adjacent to its cricket host. (**E**) *Cretacimermis longa* sp. nov. adjacent to its adult cockroach host. (**F**) *Cretacimermis longa* sp. nov. adjacent to its juvenile cockroach host. (**G**) *Cretacimermis cimicis* sp. nov. adjacent to its perforissid planthopper host. (**H**) *Cretacimeris cimicis* sp. nov. adjacent to second perforissid planthopper. Scale bars = 2.0 mm (**B**, **E**, **F**), 1.0 mm (**A**, **C**, **D**, **G**), 0.5 mm (**H**).

**Figure 2.**
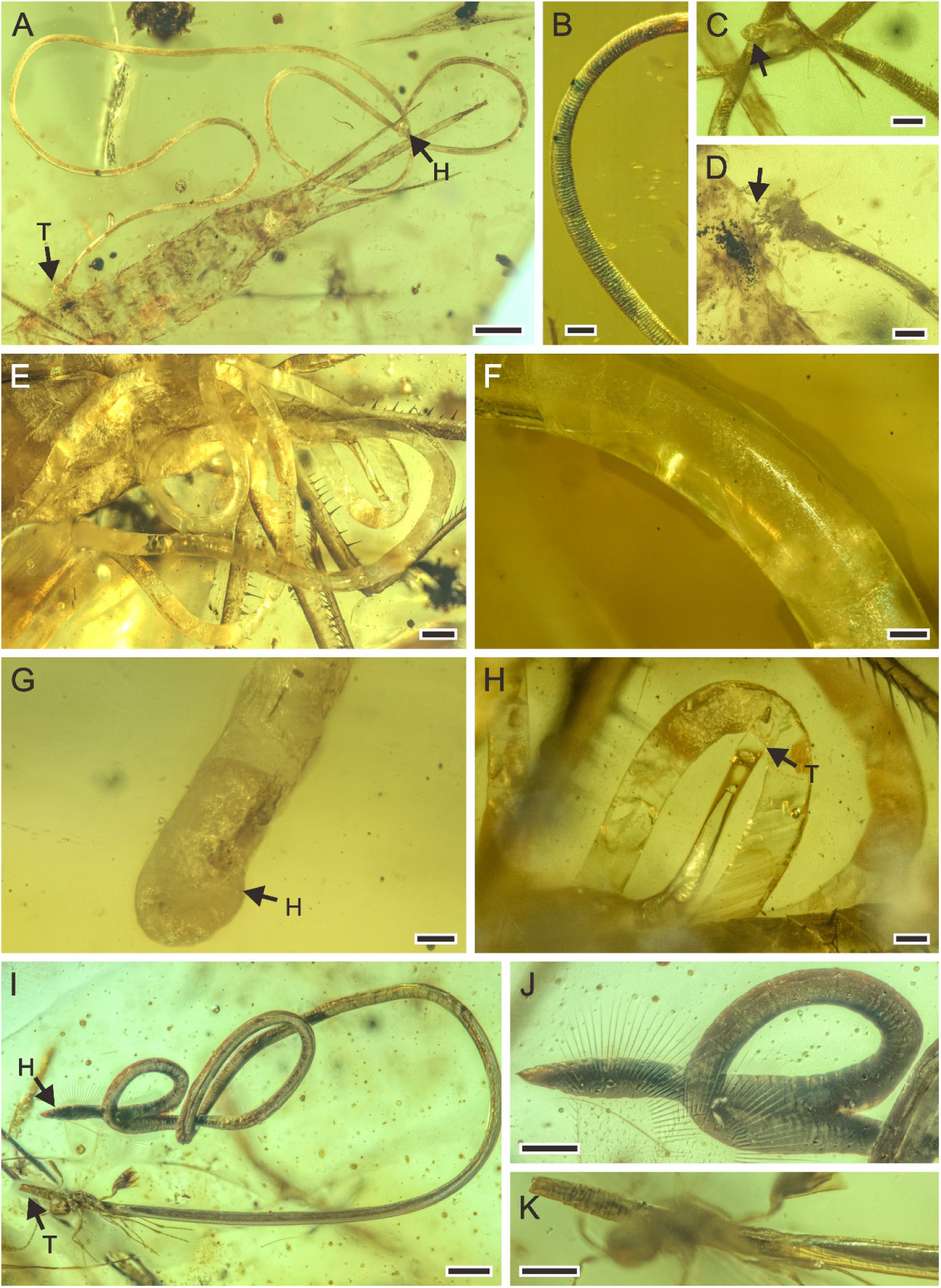
Detailed photographs of *Cretacimermis incredibilis* sp. nov., NIGP N11 (**A**–**D**), *Cretacimermis calypta* sp. nov., NIGP N08 (**E**–**H**), and *Cretacimermis adelphe* sp. nov., NIGP N18 (**I**–**K**). (**A**) habitus of *C. incredibilis*. (**B**) fine ridges in areas of body bends. (**C**) head (arrowed). (**D**) tail and the exit wound on the host (arrowed). (**E**) habitus except head part of *C. calypta*. (**F**) detail of body. (**G**) head. (**H**) tail. (**I**) habitus of upper specimen, opaque body and pointed head. (**J**) detail of head. (**K**) detail of tail. Scale bars = 0.5 mm (**A**, **E**), 0.2 mm (**H**, **I**), 0.1 mm (**B**–**D**, **F**, **G, J, K**). Abbreviation: H, head; T, tail.

urn:lsid:zoobank.org:act:

**Etymology.** The species epithet is from the Latin “incredibilis” = extraordinary.

**Type host.** Bristletail (Archaeognatha).

**Material.** Holotype. Kachin amber, rectangular piece, 18 × 7 × 4 mm, weight 0.4 g, specimen No. NIGP N11.

**Diagnosis.** Mermithid nematode parasitizing Archaeognatha from mid-Cretaceous Kachin amber.

**Description.** Body brownish, partially transparent (Figure 2A); cuticle smooth, lacking cross fibres but with fine ridges in areas of body bends (Figure 2B); trophosome evident; head rounded (Figure 2C), tail appendage not observed (Figure 2D); length 17.9 mm; greatest width 80 µm; a (length/width) = 224.

**Collective species** *Cretacimermis calypta* Luo & Poinar, sp. nov. (Figures 1B and 2E–H)

urn:lsid:zoobank.org:act:

**Etymology.** The species epithet is derived from the Greek “kalyptê” = hidden. **Type host.** A specimen of *Burmaphlebia reifi* Bechly & Poinar (Odonata: Epiophlebioptera: Epiophlebioidea: Burmaphlebiidae).

**Material.** Holotype. Kachin amber, rhomboid piece, 19 × 13 × 5 mm, weight 1.4 g, specimen No. NIGP N08.

**Diagnosis.** Mermithid nematode parasitizing Odonata from mid-Cretaceous Kachin amber.

**Description.** Body white with clear partially transparent portions (Figure 2E); cuticle lacking cross fibres (Figure 2F); trophosome fractured; head round (Figure 2G), tail bluntly rounded (Figure 2H); length 40.9 mm; greatest width 324 µm; a = 126.

**Collective species** *Cretacimermis adelphe* Luo & Poinar, sp. nov. (Figures 1C and 2I–K)

urn:lsid:zoobank.org:act:

**Etymology.** The species epithet is derived from the Greek “adelphos” = sister.

**Type host.** Earwig (Dermaptera).

**Material.** Kachin amber, cabochon, 16 × 13 × 3 mm, weight 0.4 g, specimen No. NIGP N18.

**Diagnosis.** Mermithid nematode parasitizing Dermaptera from mid-Cretaceous Kachin amber.

**Description.** Upper specimen (holotype): body dark grey, opaque, coiled (Figure 2I); cuticle smooth, lacking cross fibres; head narrowed with acute tip (Figure 2J), tail blunt (Figure 2K); length 7.4 mm, greatest width 67 µm, a = 110. Lower specimen (paratype): body dark grey, opaque, outstretched; head point-blunted, length at least 7.5 mm, greatest width 76 µm.

**Collective species** *Cretacimermis directa* Luo & Poinar, sp. nov. (Figures 1D and 3A, B)

**Figure 3.**
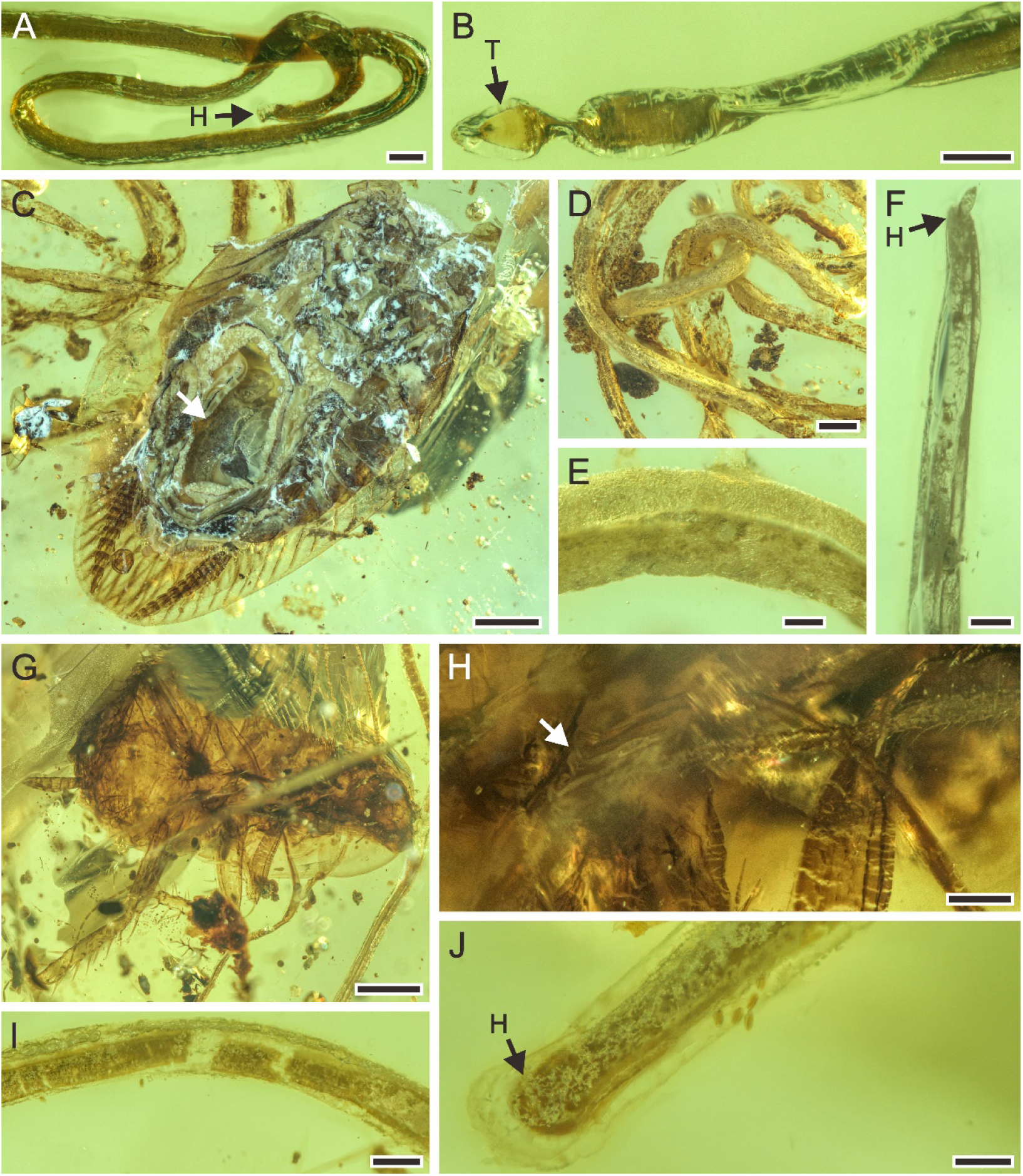
Detailed photographs of *Cretacimermis directa* sp. nov., NIGP N12 (**A**, **B**), and *Cretacimermis longa* sp. nov., NIGP N16 (**C**–**F**), NIGP N25 (**G**–**J**). (**A**) detail of head. (**B**) detail of tail. (**C**) host, an adult of Mesoblattinidae (Blattodea), note the hollow abdomen (arrowed) that probably contained the developing nematode. (**D**) detail of body. (**E**) enlarged details of body. (**F**) head. (**G**) host, a juvenile of Mesoblattinidae (Blattodea). (**H**) the terminal of nematode, note the mermithid was in the process of emerging from the host’s body (arrowed). (**I**) detail of body. (**J**) head, showing loose outer cuticle. Scale bars = 1.0 mm (**C**, **G**), 0.5 mm (**D**), 0.2 mm (**H**, **I**), 0.1 mm (**A**, **B**, **E**, **F**, **J**). Abbreviation: H, head; T, tail.

urn:lsid:zoobank.org:act:

**Etymology.** The species epithet is derived from the Latin “directis” = to/for/by the ones that are arranged in a straight line.

**Type host.** An early instar cricket nymph (Orthoptera: Ensifera: Grylloidea). **Material.** Holotype. Kachin amber, cabochon, 18.5 × 4.5 × 3 mm, weight 0.3 g, specimen No. NIGP N12.

**Diagnosis.** Mermithid nematode parasitizing Orthoptera from mid-Cretaceous Kachin amber.

**Description.** Body well preserved, elongate except for a small coil at anterior end, greyish; trophosome slightly fractured in a few areas; head rounded (Figure 3A), tail bluntly pointed (Figure 3B); length 8.8 mm; greatest width 98 µm; a = 90.

**Collective species** *Cretacimermis longa* Luo & Poinar, sp. nov. (Figures 1E, F and 3C–J)

urn:lsid:zoobank.org:act:

**Etymology.** The species epithet is from the Latin “longa” = long.

**Type host.** Cockroaches of the extinct family Mesoblattinidae (Blattodea) (Figure 3C and G).

**Material.** Kachin amber. First piece (holotype): cabochon, 20 × 13 × 2 mm, weight 0.5 g, specimen No. NIGP N16; second piece (paratype): cabochon, 35 × 24 × 6 mm, weight 5.2 g, specimen No. NIGP N25.

**Diagnosis.** Mermithid nematode parasitizing mesoblattinid cockroach from mid-Cretaceous Kachin amber.

**Description.** Nematode from adult cockroach, first piece (Figures 1E and 3C–F): body light grey, speckled, partially transparent; head narrow, tail obscured; length at least 104.3 mm, greatest width 310 µm. Nematode from juvenile cockroach, second piece (Figures 1F and 3G–J): body light to dark grey; head rounded, tail obscured; length at least 59.9 mm; greatest width 246 µm.

**Collective species** *Cretacimermis cimicis* Luo & Poinar, sp. nov. (Figures 1G, H and 4)

**Figure 4.**
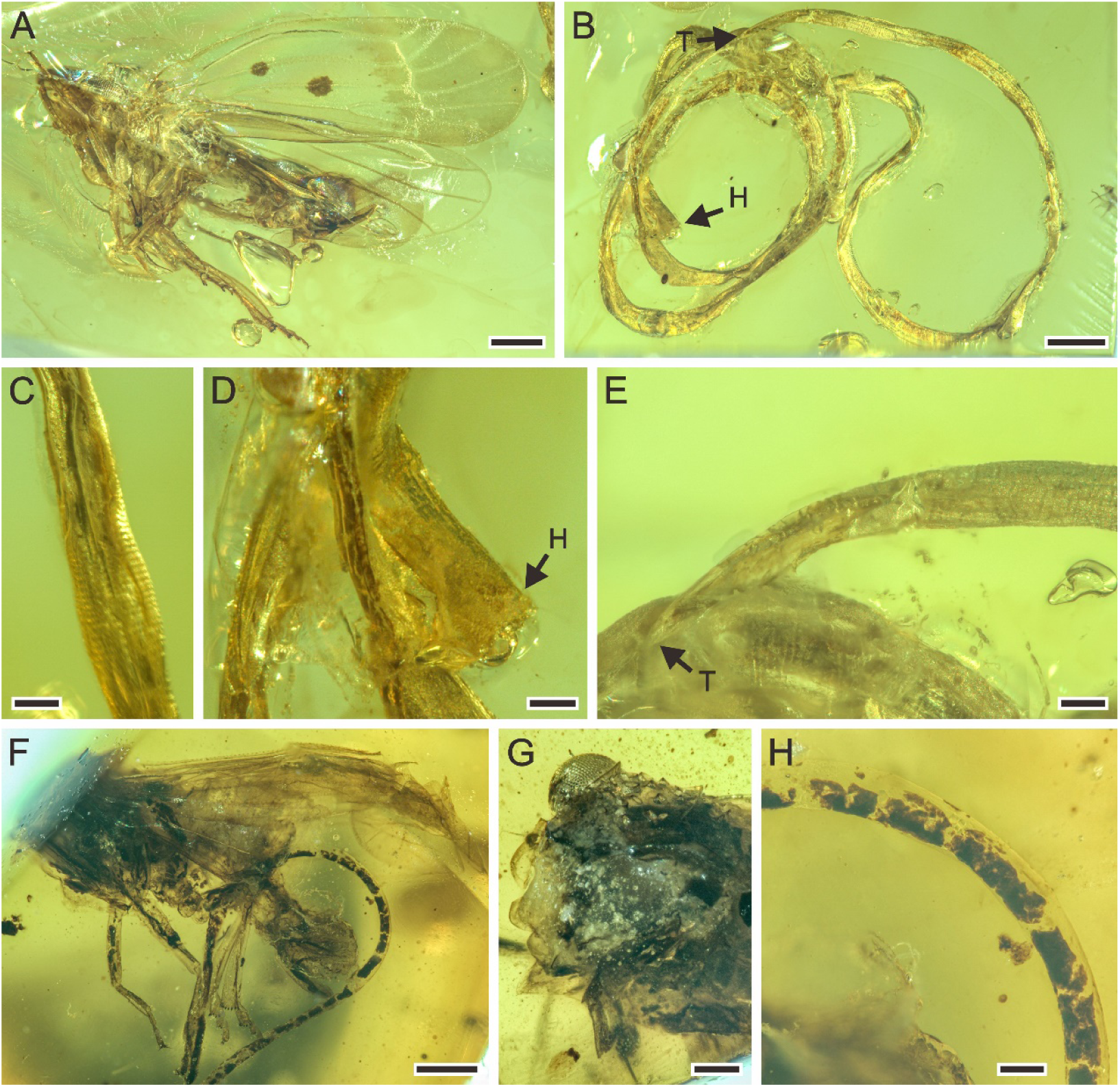
Detailed photographs of *Cretacimermis cimicis* sp. nov., NIGP N02 (**A**–**E**), and NIGP N26 (**F**–**G**). (**A**) host, Perforissidae (Hemiptera: Fulgoromorpha). (**B**) habitus of the coiled body of *C. cimicis*. (**C**) detail of body, note artefact ridges on cuticle. (**D**) head. (**E**) tail. (**F**) host, Perforissidae (Hemiptera: Fulgoromorpha). (**G**) the front view of host, indicating it is a perforissid planthopper. (**H**) detail of body, showing smooth cuticle and dark, fractured trophosome. Scale bars = 0.5 mm (**A**, **B**, **F**), 0.2 mm (**G**), 0.1 mm (**C**–**E**, **H**). Abbreviation: H, head; T, tail.

urn:lsid:zoobank.org:act:

**Etymology.** The species epithet is derived from the Latin “cimex” = bedbug.

**Type host.** Planthopper of the extinct family Perforissidae (Hemiptera: Fulgoromorpha).

**Material.** Kachin amber. First piece (holotype): trapezoid, 9 × 7 × 3 mm, weight 0.2 g, specimen No. NIGP N02; second piece (paratype): semicircular, 8 × 4 × 4 mm, weight 0.1 g, specimen No. NIGP N26.

**Diagnosis.** Mermithid nematode parasitizing perforissid planthopper from mid-Cretaceous Kachin amber.

**Description.** First piece (Figure 4A–E): body complete, mostly greyish and opaque, coiled several times; trophosome evident in some body areas; cuticle smooth, lacking cross fibres but with fine ridges in areas of body bends; head blunt; tail pointed; length 22.2 mm; greatest width 183 µm; a = 121. Second piece (Figure 4F–H): body incomplete with single coil, mostly black and opaque; trophosome fractured; cuticle smooth; head and tail missing; total length unknown; greatest width 109 µm.

**Collective species** *Cretacimermis manicapsoci* Luo & Poinar, sp. nov. (Figures 5A, B and 6A–G)

**Figure 5.**
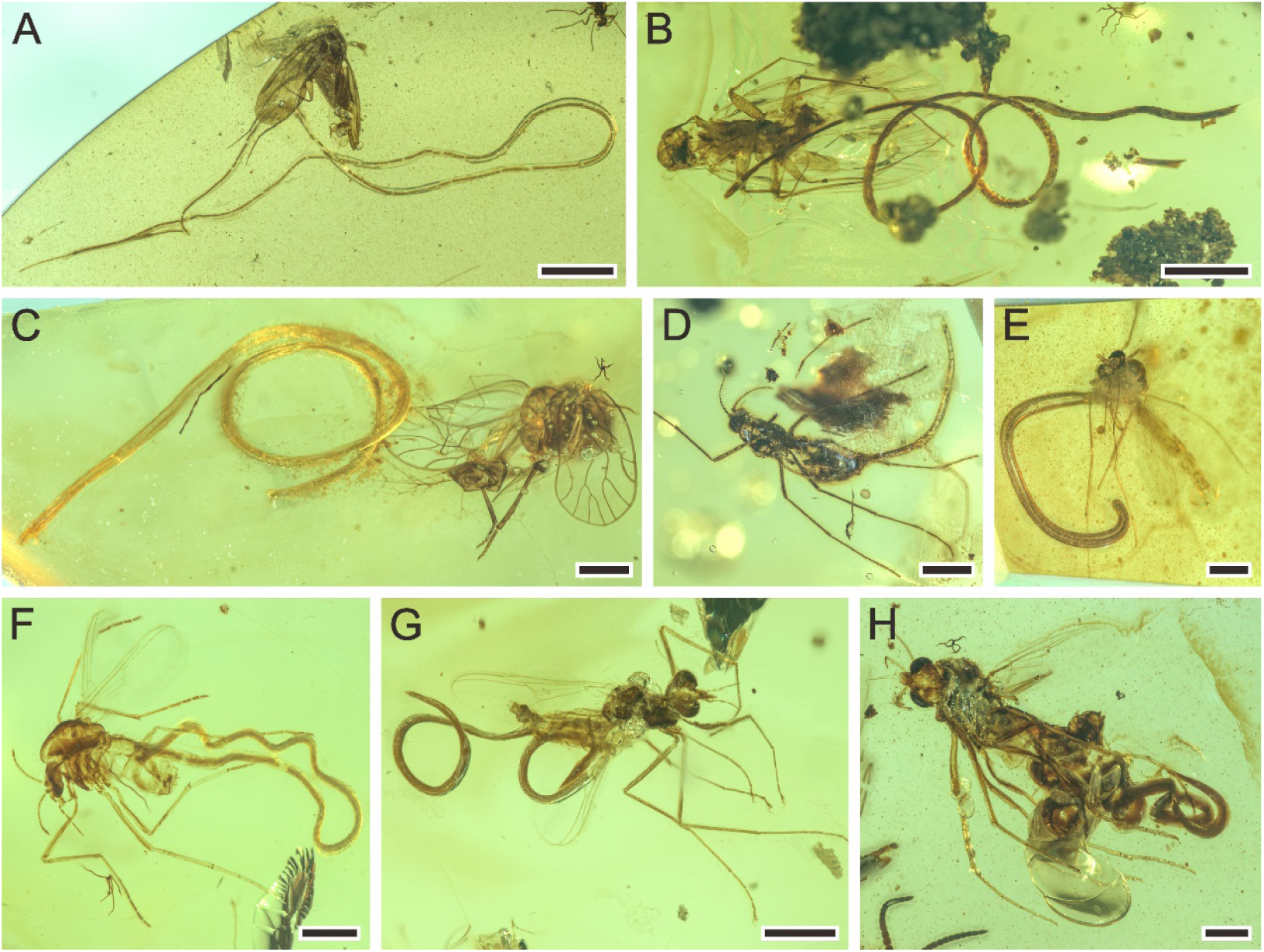
Mermithids and their insect hosts from mid-Cretaceous Kachin amber. Part II. (**A**) *Cretacimermis manicapsoci* sp. nov. adjacent to its manicapsocid barklouse host. (**B**) *Cretacimermis manicapsoci* sp. nov. adjacent to second manicapsocid barklouse host. (**C**) *Cretacimermis psoci* sp. nov. adjacent to its compsocid barklouse host. (**D**) *Cretacimermis cecidomyiae* sp. nov. emerging from its gall midge (cecidomyiid) host. (**E**–**H**) four specimens of *Cretacimermis chironomae* Poinar, 2011 emerging from their chironomid host. Scale bars = 2.0 mm (**A**), 1.0 mm (**B**), 0.5 mm (**C**–**H**).

urn:lsid:zoobank.org:act:

**Etymology.** The species epithet is derived from the family Manicapsocidae.

**Type host.** Barklouse of the family Manicapsocidae (Psocodea) (Figure 6A and E).

**Figure 6.**
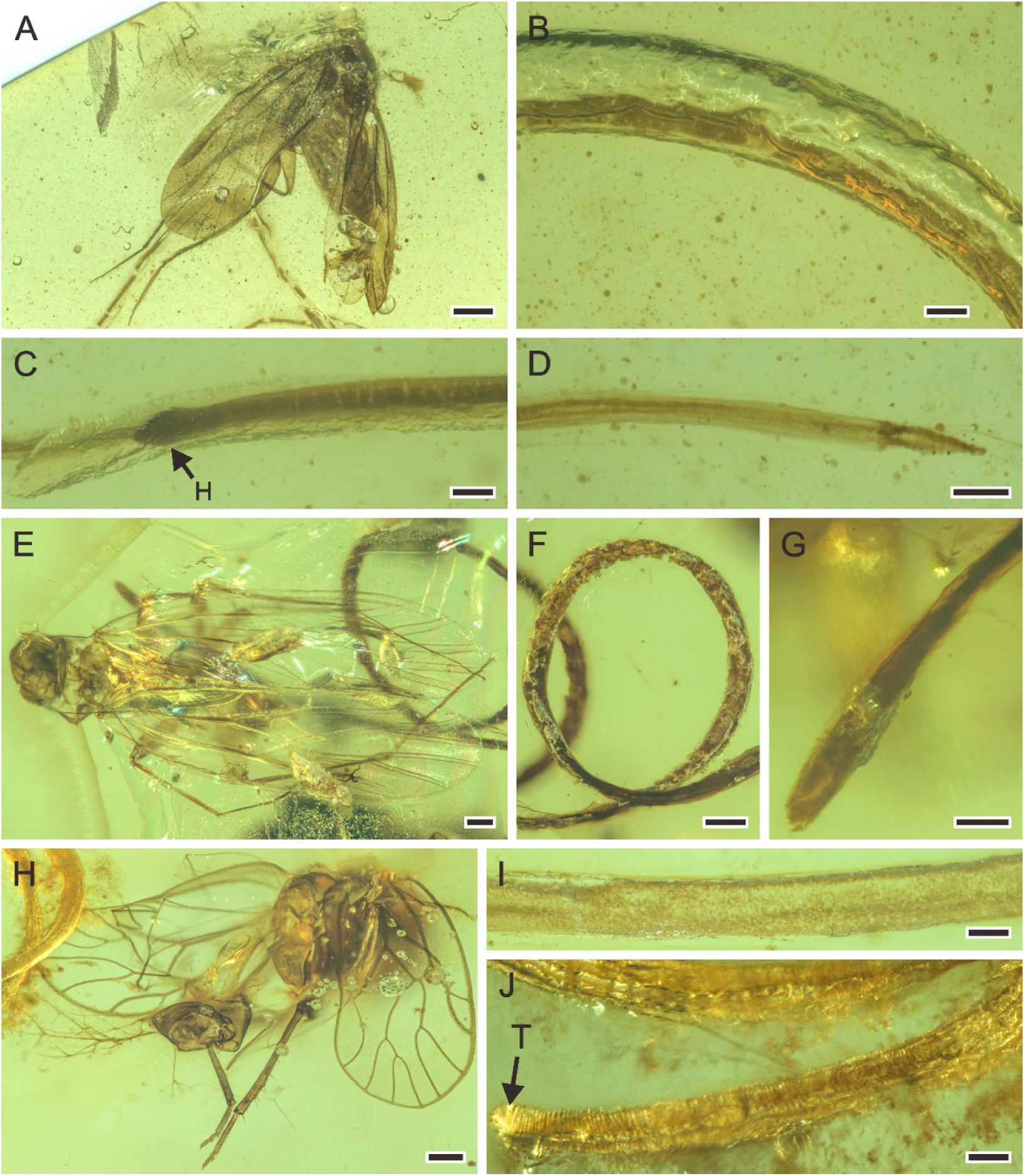
Detailed photographs of *Cretacimermis manicapsoci* sp. nov., NIGP N29 (**A**–**D**), NIGP N31 (**E**–**G**), and *Cretacimermis psoci* sp. nov., NIGP N15 (**H**–**J**). (**A**) barklouse host, Manicapsocidae (Psocoptera). (**B**) detail of body. (**C**) head. (**D**) tail. (**E**) barklouse host, Manicapsocidae. (**F**) coiled body. (**G**) head. (**H**) barklouse host, Compsocidae (Psocoptera). (**I**) detail of body. (**J**) tail, note artefactual cuticular ridges. Scale bars = 0.5 mm (**A**), 0.2 mm (**E**, **F**, **H**), 0.1 mm (**B**–**D**, **G**, **I**, **J**). Abbreviation: H, head.

**Material.** Kachin amber. First piece (holotype): cabochon, 34 × 15 × 3 mm, weight 1.3 g, specimen No. NIGP N29; second piece (paratype): cabochon, 13 × 9 × 2 mm, weight 0.1 g, specimen No. NIGP N31.

**Diagnosis.** Mermithid nematode parasitizing manicapsocid barklouse from mid-Cretaceous Kachin amber.

**Description.** First piece (Figure 6A–D): body complete, essentially a dark tube inside a clear tube, bent several times, one bend overlapping leg of host; lacking cross fibres but with fine ridges; cuticle smooth; head blunt; tail narrowed; length 34.0 mm; greatest width, 87 µm; a = 391. Second piece (Figure 6E–G): body incomplete, uneven, mostly greyish and opaque, coiled twice; trophosome evident, cuticle lacking cross fibres; head blunt, tail missing; length of remaining body 12.7 mm; greatest width 107 µm.

**Collective species** *Cretacimermis psoci* Luo & Poinar, sp. nov. (Figures 5C and 6H–J)

urn:lsid:zoobank.org:act:

**Etymology.** The species epithet is derived from New Latin “psocus” = member of psocopteran lineage.

**Type host.** Barklouse of the family Compsocidae (Psocodea) (Figure 6H).

**Material.** Holotype. Kachin amber, cabochon, 12 × 4 × 2 mm, weight 0.1 g, specimen No. NIGP N15.

**Diagnosis.** Mermithid nematode parasitizing compsocid barklouse from mid-Cretaceous Kachin amber.

**Description.** Body incomplete, tanned, partially transparent (Figure 6I); cuticle with areas of shrinkage ridges; head missing, tail rounded (Figure 6J); length 9.6 mm; greatest width 111 µm; a = 87.

**Collective species** *Cretacimermis cecidomyiae* Luo & Poinar, sp. nov. (Figures 5D and 7A–C)

**Figure 7.**
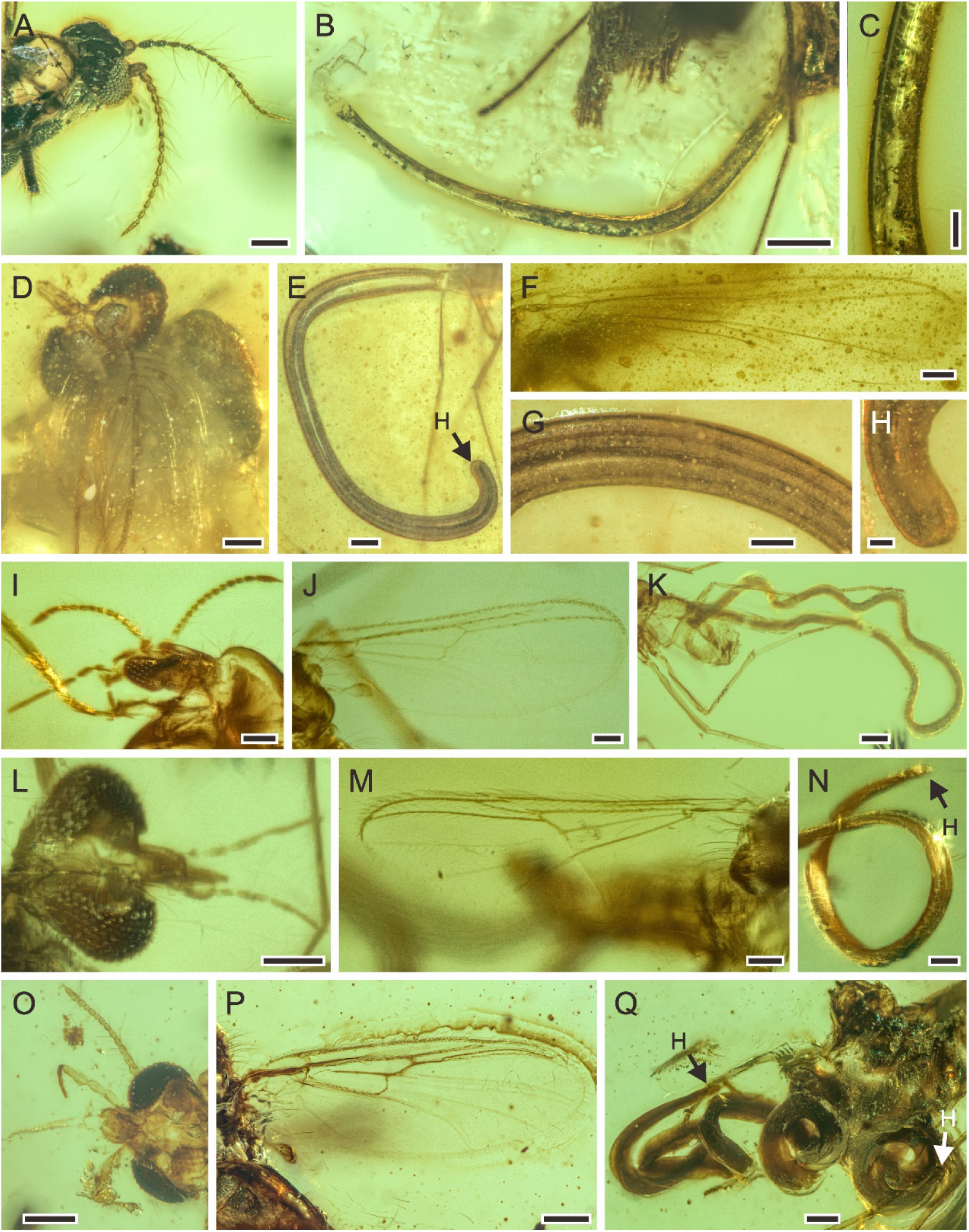
Mermithids and their Diptera hosts from mid-Cretaceous Kachin amber. (**A**–**C**), *Cretacimermis cecidomyiae* sp. nov., NIGP N09. (**A**) head of cecidomyiid host. (**B**) habitus of nematode. (**C**) detail of body. (**D**–**H**), first piece of *Cretacimermis chironomae* Poinar, 2011, NIGP N07. (**D**) detail of head of host. (**E**) overview of nematode. (**F**) forewing venation of host. (**G**) detail of body. (**H**) detail of head. (**I**– **K**), second piece of *C. chironomae*, LYD-MD-NG001. (**I**) detail of head of host. (**J**) forewing venation of host. (**K**) overview of nematode, note that a portion of the mermithid is still in the host’s abdomen. (**L**–**N**), third piece of *C. chironomae*, LYD-MD-NG002. (**L**) detail of head of host. (**M**) forewing venation of host. (**N**) detail of head and body. (**O**–**Q**), fourth piece of *C. chironomae*, NIGP N33. (**O**) detail of head of host. (**P**) forewing venation of host. (**Q**) habitus of two nematodes. Scale bars = 0.2 mm (**O**–**Q**). Scale bars = 0.2 mm (**B**, **E**, **K**, **O**–**Q**), 0.1 mm (**A**, **D**, **F**, **G**, **I**, **J**, **L**–**N**), 50 μm (**C**, **H**).

urn:lsid:zoobank.org:act:

**Etymology.** The species epithet is derived from the type genus of Cecidomyiidae “*Cecidomyia*”.

**Type host.** Gall midge of the family Cecidomyiidae (Diptera: Culicomorpha) (Figure 7A).

**Material.** Holotype. Kachin amber, cabochon, 7 × 4 × 3 mm, weight 0.1 g, specimen No. NIGP N09.

**Diagnosis.** Mermithid nematode parasitizing cecidomyiid midge from mid-Cretaceous Kachin amber.

**Description.** Body greyish with white areas; cuticle smooth, lacking cross fibres; body with dark trophosome; head missing, tail obscured; length at least 1.5 mm; greatest width 73 µm; a = at least 21 (Figure 7B and C).

**Collective species** *Cretacimermis chironomae* Poinar, 2011 (Figures 5E–H and 7D–Q)

**Type host.** A non-biting midge of the family Chironomidae (Diptera: Culicomorpha).

**Material.** Kachin amber. First piece: subtrapezoidal, 9.5 × 3.5 × 2 mm, weight 0.1 g, specimen No. NIGP N07; second piece: cabochon, 14 × 9 × 4 mm, weight 0.5 g, specimen No. LYD-MD-NG001; third piece: cabochon, 16 × 13 × 4 mm, weight 0.6 g, specimen No. LYD-MD-NG002; fourth piece: cabochon, 18 × 10 × 2 mm, weight 0.1 g, specimen No. NIGP N28

**Description.** First piece (Figure 7D–H): body greyish, uniform with little distortion; cuticle smooth, lacking cross fibres; details of trophosome clear; head rounded; length at least 3.6 mm, greatest width 193 µm. Second piece (Figure 7I–K): body well preserved, with distinct opaque trophosome; length at least 4.9 mm, greatest width 106 µm. Third piece (Figure 7L–N): body well preserved, mostly opaque due to trophosome; posterior body portion flattened and slightly twisted; head pointed; length at least 4.3 mm, greatest width 75 µm. Fourth piece (Figure 7O–Q): both specimens’ body dark brown, very wide in relation to length; cuticle smooth, lacking cross fibres; trophosome opaque; heads rounded, tails obscured; lengths not possible to attain, greatest widths 175 µm and 155 µm, respectively.

## Discussion

The sixteen new mermithids associated with their insect hosts described above include 10 insect–mermithid associations. The hosts of nine species were previously unrecorded, which triples the diversity of Cretaceous Mermithidae (from 4 to 13 species) (Supplementary Table 1). In today’s ecosystems, mermithids have been reported from a variety of arthropods, in a range of environments, and often infecting large percentages of host populations and causing mass mortality (*Poinar, 1975*, *1979; Petersen, 1985*). Despite their abundance in extant terrestrial ecosystems, mermithids are rare in the fossil record as they are not readily preserved as fossils. Twenty-two fossil mermithid species have been described from the Cenozoic with their hosts (Supplementary Table 1), mainly from Eocene Baltic amber (11 species) and Miocene Dominican amber (9 species), but only four pre-Cenozoic species associated with only two insect orders have previously been recorded (*Poinar and Buckley, 2006; Poinar and Sarto i Monteys, 2008; Poinar, 2011*, *2017*). However, according to our new records, nine insect orders are now known to have been infested by mermithid nematodes in Kachin amber and this number is even higher than that of Baltic amber (~45 Ma) and Dominican amber (~18 Ma) (six and three insect orders, respectively), despite of much longer time searching for nematodes in the latter two amber deposits (*Poinar, 2011*). Together with previously described mermithids in Kachin amber (*Poinar, 2001b; Grimaldi et al., 2002; Poinar and Buckley, 2006; Poinar, 2011*, *2017*), our results suggest that mermithid parasitism of insects was actually widespread during the mid-Cretaceous (Figure 8). Mermithids species are usually characterized by strong host specificity, they are specific to a single species or to one or two families of insects, and are almost always lethal to their hosts (*Stoffolano, 1973; Petersen, 1985*), thus our study indicates that the widespread mermithid parasitism probably already played an important role in regulating the population of insects in Cretaceous terrestrial ecosystems.

**Figure 8.**
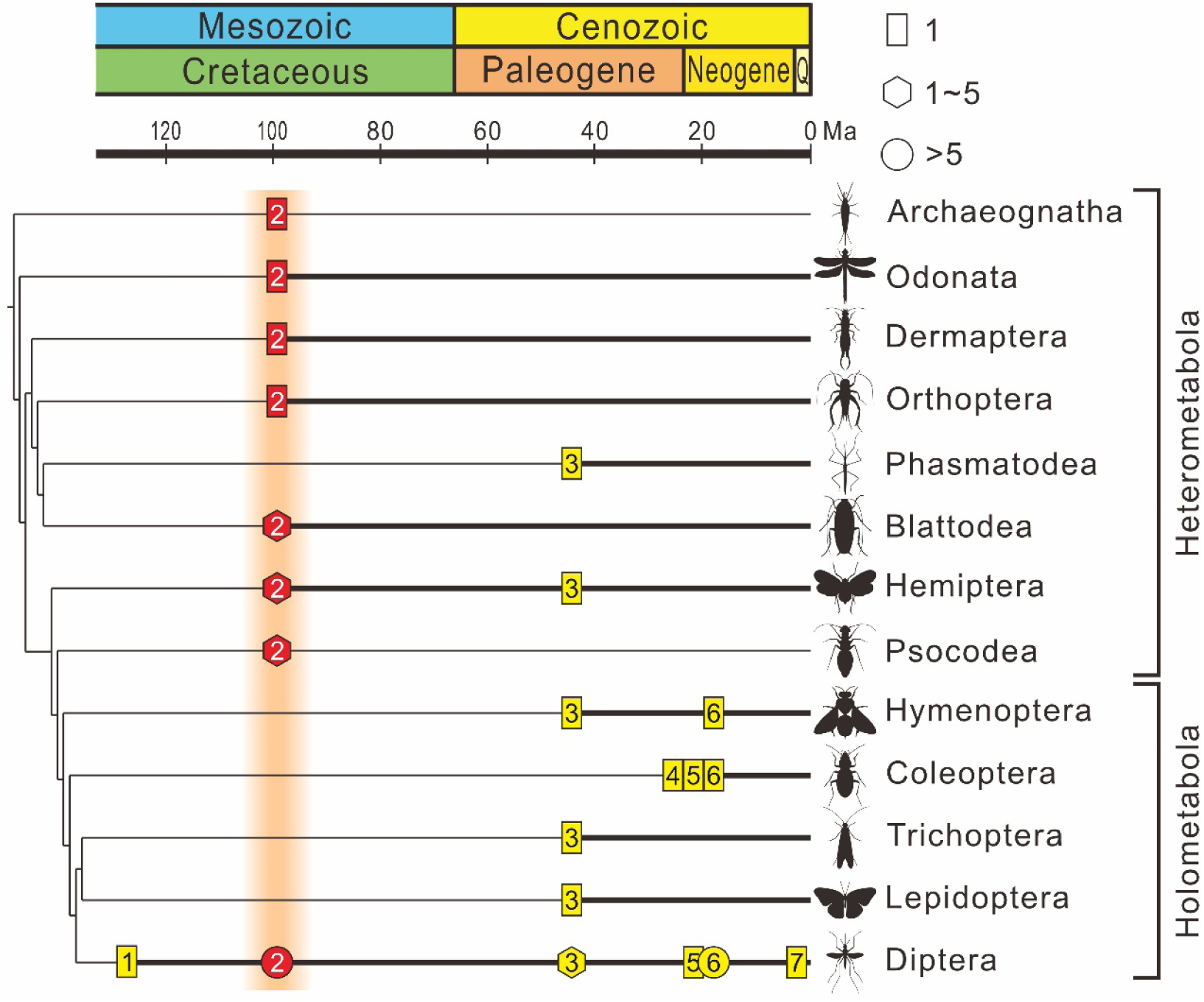
The fossil record of Mermithidae plotted on the phylogenic tree of insects. The chronogram of the insect tree is modified from Misof *et al*. (2014) (black thin line); insect orders without a fossil record of mermithid parasitism are excluded. Black thick lines indicate the presence of mermithid parasitism. Rectangles represent the fossil number of species of mermithid not exceeding one, hexagons represent the fossil number of mermithids between two and five (inclusively), and circles represent the fossil number of mermithid more than five. Yellow coloration within the symbols represents previous records, red coloration within the symbols represents records in this paper. 1 – Lebanese amber, Early Cretaceous, approximately 135 Ma; 2 – Kachin amber, mid-Cretaceous, approximately 99 Ma; 3 – Baltic amber, Eocene, approximately 45 Ma; 4 – Rhine lignite (brown coal), Oligocene/Miocene, approximately 24 Ma; 5 – Mexican amber, Early Miocene, approximately 20 Ma; 6 – Dominican amber, Miocene, approximately 18 Ma. 7 – Willershausen, Kreis Osterode, Germany, Late Pliocene, approximately 3 Ma.

Our study provides new information on fossil host–parasite associations, including three previously unknown host–mermithid associations and first fossil records of four host associations. One is *Cretacimermis incredibilis* sp. nov., which has completely exited from a bristletail (Archeognatha). Its tail end is still adjacent to what appears to be an exit wound on the host (Figure 2D), indicating a true parasitic association. There are no previous extant or extinct records of mermithids attacking bristletails (*Poinar, 1975*, *2011*). A second new mermithid–host association is barklice (Psocodea) with three different specimens parasitized by mermithids. No barklice are parasitized by mermithids today (*Poinar, 1975*), but our specimens imply that such relationship might be quite common in the mid-Cretaceous. Two members of the extinct planthopper family Perforissidae were also parasitized by mermithids, thus providing the oldest record of mermithid parasitism of planthoppers. The mermithid, *Heydenius brownii* parasitized achiliid planthoppers in Baltic amber (*Poinar, 2001a*) and this association also occurs in extant planthoppers (*Choo et al., 1989; Helden, 2008*). Furthermore, our findings are the first fossil records of mermithids parasitizing dragonflies (Odonata), earwigs (Dermaptera), crickets (Orthoptera) and cockroaches (Blattodea), four host associations predicted from extant records (*Poinar, 1975*).

Nematode body fossils are scarce and mainly known from amber, sometimes together with their hosts (*Poinar, 2011*). To explore the evolution of nematode–host relationship, we compiled nematode–host records in the three best-studied amber biotas (mid-Cretaceous Kachin amber, Eocene Baltic amber and Miocene Dominican amber) (Supplementary Table 1). Our results indicate that not only the mermithids, but also the nematodes as a whole, experienced a certain degree of host transition between the Cretaceous and Cenozoic (Figure 9). Among the insect hosts of mermithids preserved in Kachin amber, only one of the nine orders are holometabolous (i.e., insects with ‘complete’ metamorphosis), whilst it is four out of six in Baltic amber and all three insect host orders are holometabolous in Dominican amber. The situation is similar when referring to the amount of nematode parasitism. In Kachin amber, only about 40% of the hosts are Holometabola, while this percentage increases to 80% in Baltic and Dominican amber. Diptera are the most common hosts of nematodes from all three amber biotas; also, based on fossils, Diptera were the first order to be used as hosts by mermithids as depicted by fossils in Early Cretaceous Lebanese amber (*Poinar et al., 1994*). This is probably because most dipteran larvae developed in moist or aquatic environments that are particularly suitable habitats for nematodes (*Poinar, 2011*). Holometabola are the most important hosts of extant nematodes (*Poinar, 1975*), and this group dominated the insect fauna during the Cretaceous (*Labandeira and Sepkoski, 1993; Labandeira, 2005; Sohn et al., 2015; Peters et al., 2017; Zhang et al., 2018; Thomas et al., 2020; Wang et al., 2022*). Our study suggests that, except for Diptera, nematodes had not completely exploited Holometabola as hosts in the mid-Cretaceous. This suggests that heterometabolous insects (i.e., insects with ‘incomplete’ metamorphosis) were more available as hosts in the mid-Cretaceous and the widespread association between nematodes and Holometabola might be built later.

**Figure 9.**
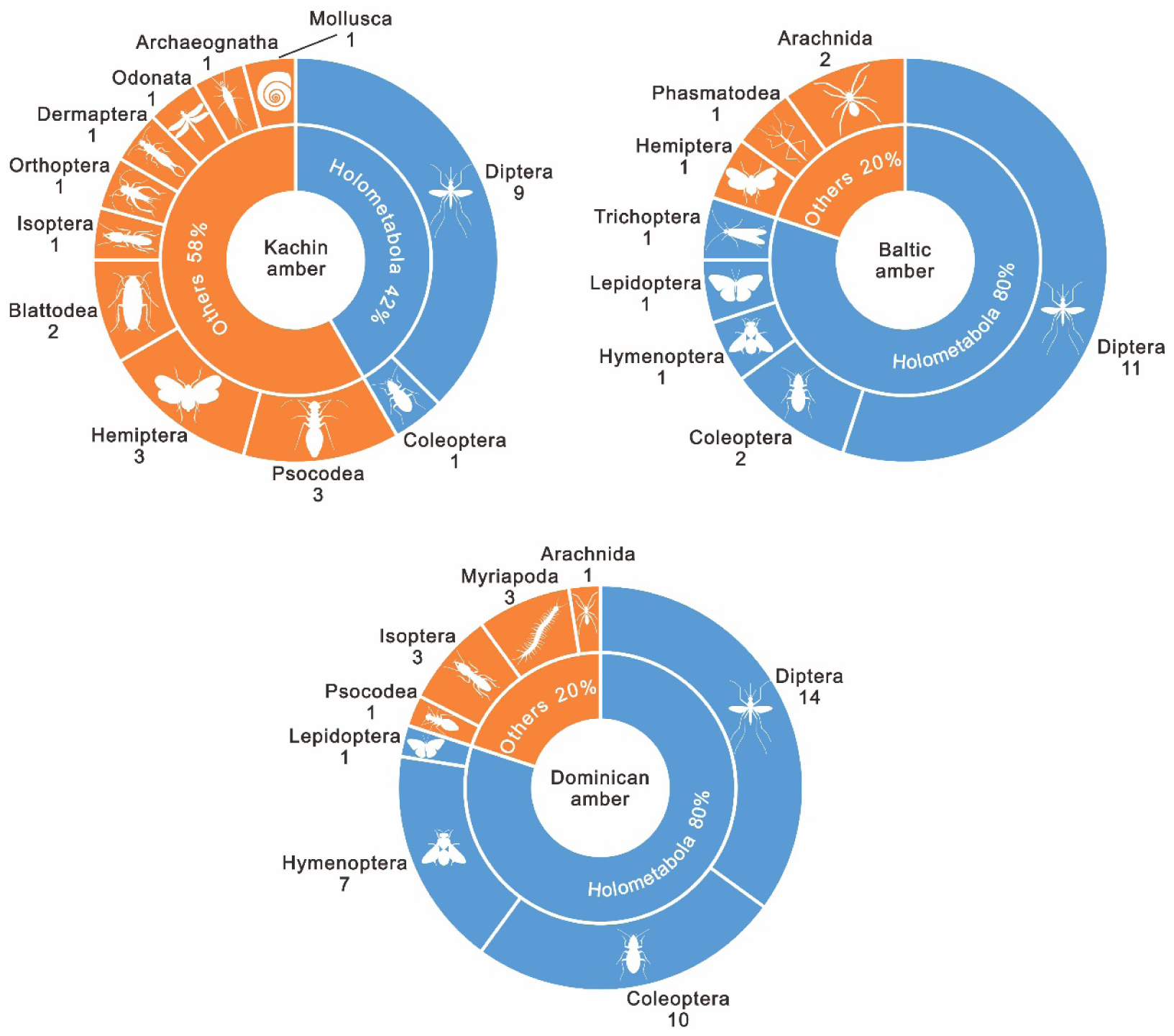
The percentage of amount of insect–nematode associations from the mid-Cretaceous Kachin amber (~99 Ma), the Eocene Baltic amber (~45 Ma) and the Miocene Dominican amber (~18 Ma). The amount of fossil species is indicated below the orders.

Finally, discovering these nematodes in mid-Cretaceous Kachin amber brings new opportunities to study the evolution of parasitism through the medium of amber. Amber is a unique form of fossilization (*Hsieh and Plotnick, 2020*). Although amber is patchily distributed in space and time, it is still especially suitable for investigating the evolution of terrestrial parasites associated with arthropods (*De Baets and Littlewood, 2015; Leung, 2017; De Baets et al., 2021; Leung, 2021*). The high diversity of mermithid nematodes during the mid-Cretaceous as shown here provides a glimpse into the structure of ancient parasitic nematode–host associations and their evolution over the past 100 million years.

## Materials and Methods

### Provenance and deposition

The specimens described here are from the Cretaceous deposits in the Hukawng Valley located southwest of Maingkhwan in Kachin State (26°20’ N, 96°36’ E) in Myanmar (*Thu and Zaw, 2017*). Radiometric U–Pb zircon dating determined the age to be 98.79 ± 0.62 Ma (*Shi et al., 2012*), a date consistent with an ammonite trapped in the amber (*Yu et al., 2019*).

Fourteen specimens (prefix NIGP) are deposited in the NIGPAS, and two specimens (prefix LYU) are deposited in Linyi University. The fossils were collected in full compliance with the laws of Myanmar and China (work on this manuscript began in early 2016). To avoid any confusion and misunderstanding, all authors declare that to their knowledge, the fossils reported in this study were not involved in armed conflict and ethnic strife in Myanmar, and were acquired prior to 2017. All specimens are permanently deposited in well-established, public museums (Supplementary Figures 1 and 2), in full compliance with the International Code of Zoological Nomenclature and the Statement of the International Palaeoentomological Society (*International Commission on Zoological Nomenclature, 1999; Szwedo et al., 2020*).

### Optical photomicrography

Observations were performed using a Zeiss Stemi 508 microscope. The photographs were taken with a Zeiss Stereo Discovery V16 microscope system in the Nanjing Institute of Geology and Palaeontology, Chinese Academy of Sciences, and measurements were taken using Zen software. Photomicrographic composites of 10 to 150 individual focal planes were digitally stacked using the software HeliconFocus 6.7.1 for a better illustration of 3D structures. Photographs were adjusted using Adobe Lightroom Classic and line drawings were prepared using CorelDraw 2019 graphic software.

## Acknowledgments

We thank Daran Zheng, Youning Su, Peter Vršanský, Adam Stroiński and Art Borkent for help with identification of the hosts of nematodes. This research was supported by the National Natural Science Foundation of China (42125201), Strategic Priority Research Program of the Chinese Academy of Sciences (XDB26000000), the Second Tibetan Plateau Scientific Expedition and Research (2019QZKK0706) and the CAS President’s International Fellowship Initiative (PIFI). This is a contribution to UNESCO-IUGS IGCP Project 679.

## Data availability

All data are available in the main text and/or the supplementary materials.

## Author contributions

Cihang Luo, Conceptualization, Methodology, Investigation, Visualization, Writing—original draft, Writing—review & editing; George O. Poinar Jr., Investigation, Visualization, Writing—original draft, Writing—review & editing; Chunpeng Xu, Investigation, Visualization, Writing—review & editing; De Zhuo, Investigation, Visualization; Edmund A. Jarzembowski, Writing—review & editing; Bo Wang, Conceptualization, Methodology, Investigation, Supervision, Writing— original draft, Writing—review & editing.

## Additional files

Supplementary Figure 1. Museum catalogue entry of Kachin amber specimens involved in this study at the Nanjing Institute of Geology and Palaeontology, Chinese Academy of Sciences.

Supplementary Figure 2. Museum catalogue entry of Kachin amber specimens involved in this study at Linyi University.

Supplementary Table 1. The fossil record of nematodes (separately uploaded).

## References

Choo HY, Kaya HK, Kim JB. 1989. *Agamermis unka* (Mermithidae) parasitism of *Nilaparvata lugens* in rice fields in Korea. Journal of Nematology 21:254–259.

De Baets K, Dentzien-Dias P, Harrison GWM, Littlewood DTJ, Parry LA, 2021. Fossil constraints on the timescale of parasitic helminth evolution, in: De Baets K, Huntley JW (Eds.), The Evolution and Fossil Record of Parasitism: Identification and Macroevolution of Parasites. Springer International Publishing, Cham, Switzerland, pp. 231–271.

De Baets K, Littlewood DTJ, 2015. Chapter one - the importance of fossils in understanding the evolution of parasites and their vectors, in: De Baets K, Littlewood DTJ (Eds.), Advances in Parasitology. Academic Press, pp. 1–51.

Grimaldi DA, Engel MS, Nascimbene PC. 2002. Fossiliferous Cretaceous amber from Myanmar (Burma): its rediscovery, biotic diversity, and paleontological significance. American Museum Novitates 2002:1–71. DOI: https://doi.org/10.1206/0003-0082(2002)361 <0001:FCAFMB>2.0.CO;2

Helden AJ. 2008. First extant records of mermithid nematode parasitism of Auchenorrhyncha in Europe. Journal of Invertebrate Pathology 99:351–353. DOI: https://doi.org/10.1016/j.jip.2008.05.005

Hsieh S, Plotnick RE. 2020. The representation of animal behaviour in the fossil record. Animal Behaviour 169:65–80. DOI: https://doi.org/10.1016/j.anbehav.2020.09.010

International Commission on Zoological Nomenclature. 1999. International Code of Zoological Nomenclature (4th ed.). International Trust for Zoological Nomenclature, London, UK.

Labandeira CC, 2005. Fossil history and evolutionary ecology of Diptera and their associations with plants, in: Yeates DK, Wiegmann BM (Eds.), The Evolutionary Biology of Flies. Columbia University Press, New York, the USA, pp. 217–273.

Labandeira CC, Sepkoski JJ, Jr. 1993. Insect diversity in the fossil record. Science 261:310–315. DOI: https://doi.org/10.1126/science.11536548

Leung TLF. 2017. Fossils of parasites: what can the fossil record tell us about the evolution of parasitism? Biological Reviews 92:410–430. DOI: https://doi.org/10.1111/brv.12238

Leung TLF, 2021. Parasites of fossil vertebrates: what we know and what can we expect from the fossil record?, in: De Baets K, Huntley JW (Eds.), The Evolution and Fossil Record of Parasitism: Identification and Macroevolution of Parasites. Springer Nature, Cham, Switzerland, pp. 1–28.

Lorenzen S, 1994. The phylogenetic systematics of freeliving nematodes. The Ray Society.

Misof B, Liu S, Meusemann K, Peters RS, Donath A, Mayer C, Frandsen PB, Ware J, Flouri T, Beutel RG, Niehuis O, Petersen M, Izquierdo-Carrasco F, Wappler T, Rust J, Aberer AJ, Aspöck U, Aspöck H, Bartel D, Blanke A, Berger S, Böhm A, Buckley TR, Calcott B, Chen J, Friedrich F, Fukui M, Fujita M, Greve C, Grobe P, Gu S, Huang Y, Jermiin LS, Kawahara AY, Krogmann L, Kubiak M, Lanfear R, Letsch H, Li Y, Li Z, Li J, Lu H, Machida R, Mashimo Y, Kapli P, McKenna DD, Meng G, Nakagaki Y, Navarrete-Heredia JL, Ott M, Ou Y, Pass G, Podsiadlowski L, Pohl H, von Reumont BM, Schütte K, Sekiya K, Shimizu S, Slipinski A, Stamatakis A, Song W, Su X, Szucsich NU, Tan M, Tan X, Tang M, Tang J, Timelthaler G, Tomizuka S, Trautwein M, Tong X, Uchifune T, Walzl MG, Wiegmann BM, Wilbrandt J, Wipfler B, Wong TKF, Wu Q, Wu G, Xie Y, Yang S, Yang Q, Yeates DK, Yoshizawa K, Zhang Q, Zhang R, Zhang W, Zhang Y, Zhao J, Zhou C, Zhou L, Ziesmann T, Zou S, Li Y, Xu X, Zhang Y, Yang H, Wang J, Wang J, Kjer KM, Zhou X. 2014. Phylogenomics resolves the timing and pattern of insect evolution. Science 346:763–767. DOI: https://doi.org/10.1126/science.1257570

Nickle WR. 1972. A contribution to our knowledge of the Mermithidae (Nematoda). Journal of Nematology 4:113–146.

Peters RS, Krogmann L, Mayer C, Donath A, Gunkel S, Meusemann K, Kozlov A, Podsiadlowski L, Petersen M, Lanfear R, Diez PA, Heraty J, Kjer KM, Klopfstein S, Meier R, Polidori C, Schmitt T, Liu S, Zhou X, Wappler T, Rust J, Misof B, Niehuis O. 2017. Evolutionary history of the Hymenoptera. Current Biology 27:1013–1018. DOI: https://doi.org/10.1016/j.cub.2017.01.027

Petersen JJ, 1985. Nematodes as biological control agents: Part I. Mermithidae, in: Baker JR, Muller R (Eds.), Advances in Parasitology. Academic Press, pp. 307–344.

Poinar GO, Jr., 1975. Entomogenous nematodes: a manual and host list of insect-nematode associations. E. J. Brill, Leiden, The Netherlands.

Poinar GO, Jr., 1979. Nematodes for biological control of insects, 1st ed. CRC Press, Boca Raton, USA.

Poinar GO, Jr., 1983. The natural history of Nematodes. Prentice Hall, Englewood Cliffs.

Poinar GO, Jr. 2001a. *Heydenius brownii* sp. n. (Nematoda: Mermithidae) parasitising a planthopper (Homoptera: Achilidae) in Baltic amber. Nematology 3:753–757. DOI: https://doi.org/10.1163/156854101753625263

Poinar GO, Jr., 2001b. Nematoda and Nematomorpha, in: Thorp JH, Covich AP (Eds.), Ecology and Classification of North American Freshwater Invertebrates, Second Edition ed. Academic Press, New York, pp. 255–295.

Poinar GO, Jr., 2011. The evolutionary history of nematodes: as revealed in stone, amber and mummies. Brill, Amersfoort, the Netherlands.

Poinar GO, Jr., 2015. Chapter 14 - Phylum Nemata, in: Thorp JH, Rogers DC (Eds.), Thorp and Covich’s Freshwater Invertebrates (Fourth Edition). Academic Press, Boston, pp. 273–302.

Poinar GO, Jr. 2017. A mermithid nematode, Cretacimermis aphidophilus sp. n. (Nematoda: Mermithidae), parasitising an aphid (Hemiptera: Burmitaphididae) in Myanmar amber: a 100 million year association. Nematology 19:509–513. DOI: https://doi.org/10.1163/15685411-00003063

Poinar GO, Jr., Acra A, Acra F. 1994. Earliest fossil nematode (Mermithidae) in Cretaceous Lebanese amber. Fundamental and Applied Nematology 17:475–477.

Poinar GO, Jr., Buckley R. 2006. Nematode (Nematoda: Mermithidae) and hairworm (Nematomorpha: Chordodidae) parasites in Early Cretaceous amber. Journal of Invertebrate Pathology 93:36–41. DOI: https://doi.org/10.1016/j.jip.2006.04.006

Poinar GO, Jr., Kerp H, Hass H. 2008. *Palaeonema phyticum gen*. n., sp. n. (Nematoda: Palaeonematidae fam. n.), a Devonian nematode associated with early land plants. Nematology 10:9–14. DOI: https://doi.org/10.1163/156854108783360159

Poinar GO, Jr., Otieno WA. 1974. Evidence of four molts in the Mermithidae. Nematologica 20:370. DOI: https://doi.org/https://doi.org/10.1163/187529274X00456

Poinar GO, Jr., Sarto i Monteys V. 2008. Mermithids (Nematoda: Mermithidae) of biting midges (Diptera: Ceratopogonidae): *Heleidomermis cataloniensis* n. sp. from Culicoides circumscriptus Kieffer in Spain and a species of *Cretacimermis* Poinar, 2001 from a ceratopogonid in Burmese amber. Systematic Parasitology 69:13–21. DOI: https://doi.org/10.1007/s11230-007-9091-9

Shi G, Grimaldi DA, Harlow GE, Wang J, Wang J, Yang M, Lei W, Li Q, Li X. 2012. Age constraint on Burmese amber based on U–Pb dating of zircons. Cretaceous Research 37:155–163. DOI: https://doi.org/10.1016/j.cretres.2012.03.014

Sohn J-C, Labandeira CC, Davis DR. 2015. The fossil record and taphonomy of butterflies and moths (Insecta, Lepidoptera): implications for evolutionary diversity and divergence-time estimates. BMC Evolutionary Biology 15:12. DOI: https://doi.org/10.1186/s12862-015-0290-8

Stoffolano JG, Jr. 1973. Host specificity of entomophilic nematodes—A review. Experimental Parasitology 33:263–284. DOI: https://doi.org/10.1016/0014-4894(73)90033-7

Szwedo J, Wang B, Soszynska-Maj A, Azar D, Ross A. 2020. International Palaeoentomological Society Statement. Palaeoentomology 3:221–222. DOI: https://doi.org/10.11646/palaeoentomology.3.3.1

Thomas JA, Frandsen PB, Prendini E, Zhou X, Holzenthal RW. 2020. A multigene phylogeny and timeline for Trichoptera (Insecta). Systematic Entomology 45:670–686. DOI: https://doi.org/10.1111/syen.12422

Thu K, Zaw K, 2017. Chapter 23 - Gem deposits of Myanmar, in: Barber AJ, Zaw K, Crow MJ (Eds.), Myanmar: Geology, Resources and Tectonics. The Geological Society, London, UK, pp. 497–529.

van den Hoogen J, Geisen S, Routh D, Ferris H, Traunspurger W, Wardle DA, de Goede RGM, Adams BJ, Ahmad W, Andriuzzi WS, Bardgett RD, Bonkowski M, Campos-Herrera R, Cares JE, Caruso T, de Brito Caixeta L, Chen X, Costa SR, Creamer R, Mauro da Cunha Castro J, Dam M, Djigal D, Escuer M, Griffiths BS, Gutiérrez C, Hohberg K, Kalinkina D, Kardol P, Kergunteuil A, Korthals G, Krashevska V, Kudrin AA, Li Q, Liang W, Magilton M, Marais M, Martín JAR, Matveeva E, Mayad EH, Mulder C, Mullin P, Neilson R, Nguyen TAD, Nielsen UN, Okada H, Rius JEP, Pan K, Peneva V, Pellissier L, Carlos Pereira da Silva J, Pitteloud C, Powers TO, Powers K, Quist CW, Rasmann S, Moreno SS, Scheu S, Setälä H, Sushchuk A, Tiunov AV, Trap J, van der Putten W, Vestergård M, Villenave C, Waeyenberge L, Wall DH, Wilschut R, Wright DG, Yang J-i, Crowther TW. 2019. Soil nematode abundance and functional group composition at a global scale. Nature 572:194–198. DOI: https://doi.org/10.1038/s41586-019-1418-6

Wang B, Xu C, Jarzembowski EA. 2022. Ecological radiations of insects in the Mesozoic. Trends in Ecology & Evolution 37:529–540. DOI: https://doi.org/10.1016/j.tree.2022.02.007

Yeates GW, Ferris H, Moens T, Van Der Putten WH, 2009. The role of nematodes in ecosystems, in: Wilson MJ, Kakouli-Duarte T (Eds.), Nematodes as environmental indicators. CAB International, Wallingford, UK, pp. 1–44.

Yu T, Kelly R, Mu L, Ross A, Kennedy J, Broly P, Xia F, Zhang H, Wang B, Dilcher D. 2019. An ammonite trapped in Burmese amber. Proceedings of the National Academy of Sciences 116:11345–11350. DOI: https://doi.org/10.1073/pnas.1821292116

Zhang S-Q, Che L-H, Li Y, Dan L, Pang H, Ślipiński A, Zhang P. 2018. Evolutionary history of Coleoptera revealed by extensive sampling of genes and species. Nature Communications 9:205. DOI: https://doi.org/10.1038/s41467-017-02644-4

Zhang Y, Li S, Li H, Wang R, Zhang K-Q, Xu J. 2020. Fungi–nematode interactions: diversity, ecology, and biocontrol prospects in agriculture. Journal of Fungi 6:206. DOI: https://doi.org/10.3390/jof6040206

